# Cytokinin perception in potato: New features of canonic players

**DOI:** 10.1101/269266

**Authors:** Sergey N. Lomin, Yulia A. Myakushina, Oksana O. Kolachevskaya, Irina A. Getman, Dmitry V. Arkhipov, Ekaterina M. Savelieva, Dmitry I. Osolodkin, Georgy A. Romanov

## Abstract

Potato is the most economically important non-cereal food crop. Tuber formation in potato is regulated by phytohormones, cytokinins (CKs) in particular. The present work was aimed to study CK signal perception in potato. The sequenced potato genome of doubled monoploid Phureja was used for bioinformatic analysis and as a tool for identification of putative CK receptors from autotetraploid potato cv. Désirée. All basic elements of multistep phosphorelay (MSP) required for CK signal transduction were identified in Phureja genome, including three genes orthologous to three CK receptor genes (*AHK 2-4*) of Arabidopsis. As distinct from Phureja, autotetraploid potato contains at least two allelic isoforms of each receptor type. Putative receptor genes from Désirée plants were cloned, sequenced and expressed, and main characteristics of encoded proteins, firstly their consensus motifs, structure models, ligand-binding properties, and the ability to transmit CK signal, were determined. In all studied aspects the predicted sensor histidine kinases met the requirements for genuine CK receptors. Expression of potato CK receptors was found to be organ-specific and sensitive to growth conditions, particularly to sucrose content. Our results provide a solid basis for further in-depth study of CK signaling system and biotechnological improvement of potato.

## Introduction

Potato is a widespread practically important crop, its tuber formation is controlled by phytohormones (reviewed in Aksenova *et al*., 2012, 2014). Previous studies have shown that cytokinins (CKs) and auxins can accelerate and enhance potato tuber formation (Aksenova *et al*., 2000; Romanov *et al*., 2000; Roumeliotis *et al*., 2012; Kolachevskaya *et al*., 2015, 2017; Wang *et al*., 2018). In non-tuberizing plants (tobacco, tomato), increased doses of active CKs stimulate morphogenesis, in many aspects resembling tuber formation (Guivarc’h *et al*., 2002; Eviatar-Ribak *et al*., 2013). CK signaling is also involved in the formation of nodules on the roots of legumes (reviewed in Frugier *et al*., 2008; Miri *et al*., 2016). CKs largely determine the nature of source-sink relationships in the whole plant, enhancing the attracting ability of the tubers (Abelenda and Prat, 2013). Elevated doses of CKs affect the overall architectonics of potato plants, suppressing the root development (Aksenova *et al*., 2000). In addition, CKs participate in plant defense against biotic and abiotic adverse factors (Zwack and Rashotte, 2015; Brütting *et al*., 2017; Thu *et al*., 2017). All the above indicates the important role of CKs in both the formation of tubers and the general development and resistance of potato plants.

The molecular mechanism of CK action on a plant cell has been established using mainly the Arabidopsis model (reviewed in Hutchison and Kieber, 2002, Hwang *et al*., 2002; Kakimoto, 2003; Heyl and Schmülling, 2003; Sakakibara, 2006; Müller and Sheen, 2007). This mechanism is based on multistep phosphorelay (MSP) and uses three protein species to bring the CK signal up to the primary response genes: (i) transmembrane catalytic receptors with histidine kinase activity, (ii) mobile phosphotransmitters circulating between the cytoplasm and nucleus, and (iii) nuclear transcription factors, B-type response regulators. Other proteins (CRFs, pseudophosphotransmitters, A-type response regulators) affect the intensity of the CK signaling through the main transmission pathway (Kieber and Schaller, 2014, 2018).

Receptors are key factors in the perception and transduction of hormonal signals. In the case of CKs, receptors are sensory hybrid histidine kinases largely homologous to bacterial sensory histidine kinases, members of two-component signal transduction system. Known CK receptors are multidomain proteins located mainly in ER membranes (Caesar *et al*., 2011; Lomin *et al*., 2011, 2018; Wulfetange *et al*., 2011; Daudu *et al*., 2017; Ding *et al*., 2017) with N-terminal hormone-binding sensory module localized in the ER lumen and the central and C-terminal catalytic domains protruding in the cytosol (Steklov *et al*., 2013; Lomin *et al*., 2018). Until now, CK receptors have been studied in a few vascular plant species, primarily and most detailed in Arabidopsis and maize (Kakimoto, 2003; Yonekura-Sakakibara *et al*., 2004; Romanov *et al*., 2006; Lomin *et al*., 2011, 2012, 2015; 2018; Stolz *et al*., 2011; Heyl *et al*., 2012; Steklov *et al*., 2013; Wang *et al*., 2014). In recent years, CK receptor studies have been extended to new species including rice (Choi *et al*., 2012; Ding *et al*., 2017), *Lotus japonicus* (Held *et al*., 2014), *Medicago truncatula* (Laffont *et al*., 2015; Boivin *et al*., 2016), oilseed rape (Kuderová *et al*., 2015), *Nicotiana attenuata* (Schäfer *et al*., 2015), and apple (Daudu *et al*., 2017). These studies have demonstrated that the CK perception apparatus in some aspects is species-specific. Potato differs from most plant species by its ability to form tubers. This process, sensitive to various cues including CKs, makes the study of CK receptors of potato especially intriguing. So far, to our knowledge, there have been no scientific reports on such studies.

In this paper, we have examined potato CK receptors of a homozygous doubled monoploid Phureja (DM1-3 516 R44) whose genome was sequenced several years ago (Potato Genome Sequencing Consortium, 2011). Cloning and expression of receptor encoding genes were conducted using the commercial autotetraploid potato cv. Désirée. The presence of all necessary MSP elements in potato was demonstrated and main characteristics of CHASE domain-containing CK receptors, primarily their consensus motifs, 3D structure models, ligand-binding properties, and the ability to transmit the signal by MSP were ascertained. In contrast to the Phureja monoploid, distinct alleles for each of the three main forms of receptors were found in the Désirée potato. Expression of CK receptor genes was shown to be organ-specific and affected by sucrose. The obtained results might serve as a framework for new biotechnological approaches in improving potato productivity and stress resistance.

## Materials and methods

### Sequence analysis

Nucleotide/polypeptide sequences of CK receptors and other proteins related to the CK signaling were retrieved from databases NCBI (National Center for Biotechnology Information, http://www.ncbi.nlm.nih.gov), Phytozome 11 (https://phytozome.jgi.doe.gov/pz/portal.html), MSU Rice Genome Annotation Project Release 7 (http://rice.plantbiology.msu.edu/) and congenie.org (http://congenie.org/) using the BLASTP tool and *AHK2* (AT5G35750), *AHK3* (AT1G27320), *AHK4* (AT2G01830) and other CK-related genes of *Arabidopsis thaliana* as templates. Domain structure of proteins was determined with PROSITE (http://prosite.expasy.org/). Transmembrane domains were determined using MESSA service (http://prodata.swmed.edu/MESSA/MESSA.cgi) (Cong and Grishin, 2012). Domain visualization was performed using the MyDomains – Image Creator service (http://prosite.expasy.org/mydomains/).

Phylogenetic analysis was performed using the MEGA6.0 (Tamura *et al*., 2013). Alignment of nucleotide sequences (CDS, codon mode) was performed by ClustalW algorithm. Method of maximum likelihood was employed for phylogenetic reconstruction. The search for key amino acids (aa) in receptor domains by alignment and visualization of protein sequences was carried out in Clustal X2.1 (Larkin *et al*., 2007) and Jalview (Clamp *et al*., 2004), respectively.

### Homology modeling

Search of templates for homology modeling was performed at SWISS-MODEL web-service (https://swissmodel.expasy.org/) (Biasini *et al*., 2014). Modeling of potato (*Solanum tuberosum* L.) protein structures was accomplished in Modeller 9.19 (https://salilab.org/modeller/) (Sali and Blundell, 1993) using *automodel* class for comparative modeling. For each protein, 200 models were built, and the best model was selected according to DOPE score value (Shen and Sali, 2006) as determined by Modeller. Templates for modeling and respective references (Müller-Dieckmann *et al*., 1999; Hothorn *et al*., 2011; Pekárová *et al*., 2011; Bauer *et al*., 2013; Mayerhofer *et al*., 2015; Dubey *et al*., 2016) are listed in Supplementary Table 1. After adding hydrogen atoms, models were energy minimized in USCF Chimera 1.12 (http://www.cgl.ucsf.edu/chimera/) (Pettersen *et al*., 2004) using AMBER ff14SB force field (Maier *et al*., 2015) with 300 steps of steepest descent and 300 steps of conjugate gradient optimization; step size was 0.02 Å in both cases. Stereochemical quality of the models was assessed with ProCheck (Laskowski *et al*., 1993) implemented in PDBsum Web service (www.ebi.ac.uk/pdbsum), ProSA-web (https://prosa.services.came.sbg.ac.at/prosa.php) (Wiederstein and Sippl, 2007) and QMEAN server (https://swissmodel.expasy.org/qmean/help) (Benkert *et al*., 2009). Visualization and superposition of the models were accomplished with UCSF Chimera.

**Table 1.**
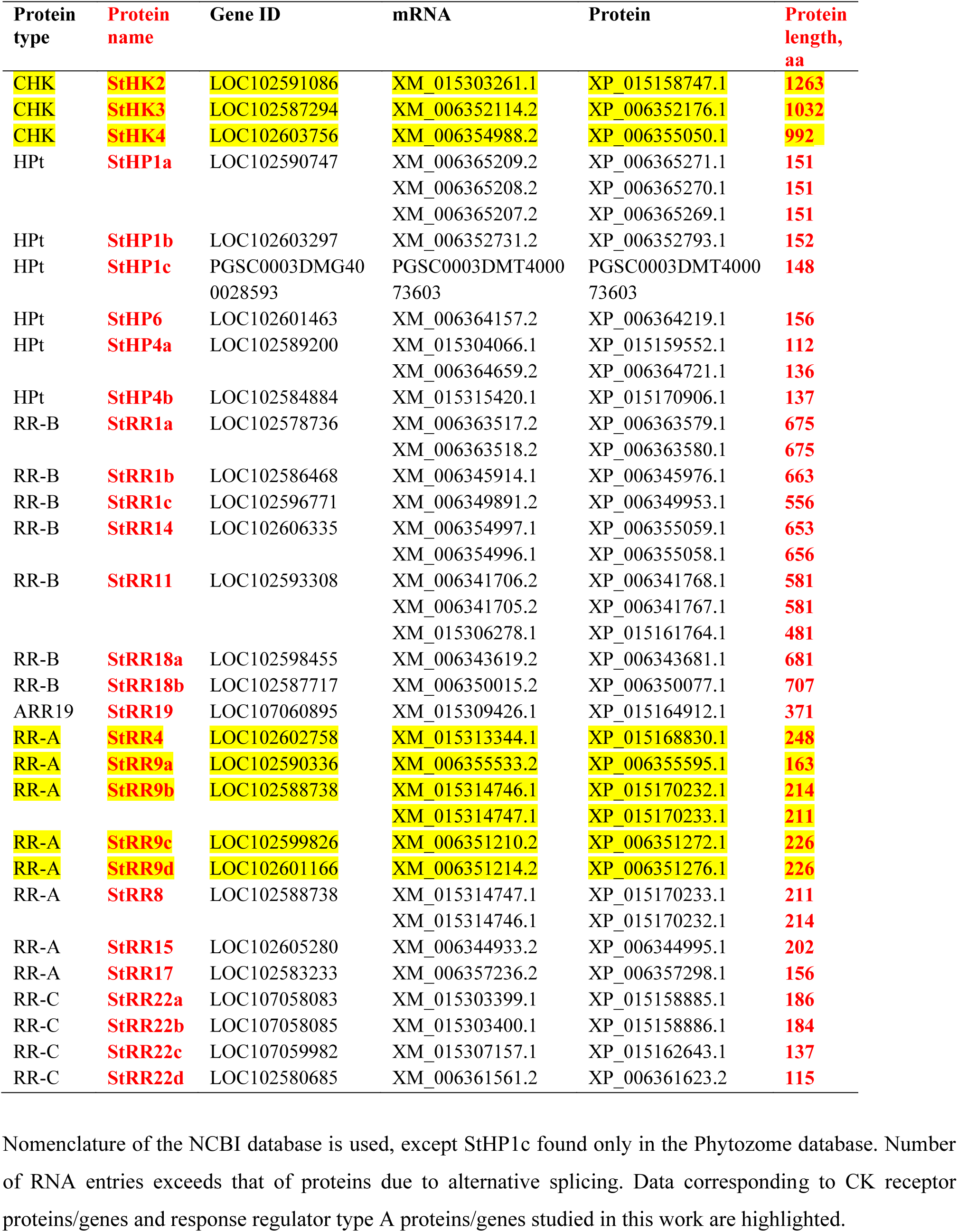
Proteins and genes predictably related to CK signaling system of potato

### Promoter analysis

Promoter regions of *Arabidopsis thaliana* CK receptor genes (*AHK2*, *AHK3* and *AHK4*) were obtained from TAIR database (https://www.arabidopsis.org). Identification of promoter regions of CK receptor genes (*StHK2*, *StHK3* and *StHK4*) of potato was performed using Phytozome 11 and NCBI databases. DNA sequence of 1000 nucleotides long upstream the gene transcription start was taken as a promoter region. The search for *cis*-regulatory elements in promoters of studied genes was carried out using PLACE (http://www.dna.affrc.go.jp/htdocs/PLACE/) and PlantCARE (http://bioinformatics.psb.ugent.be/webtools/plantcare/html/) programs.

### Receptor cloning

Experiments were performed with autotetraploid potato (*Solanum tuberosum* L.) plantlets of Désirée variety. Plants were propagated by *in vitro* cloning on Murasige-Skoog agarose medium supplemented with 1.5% sucrose, at 20 °C and 16 h photoperiod in a controlled climate chamber with luminescent white light illumination (Kolachevskaya *et al*., 2015, 2017). Total RNA was isolated from single potato shoots and treated with RNase-free DNase I (Thermo Scientific). Reverse transcription was performed with RevertAid ™ according to the manufacturer’s instructions (Thermo Scientific). Total DNA was isolated from shoots of individual plants using the CTAB method. The resulting cDNA and total DNA were used to amplify genes encoding predicted potato CK receptors with high-precision Phusion High-Fidelity DNA polymerase (Thermo Scientific). The primer design was performed to amplify the full-length and truncated (sensory modules with flanking transmembrane helices) CDS of CK receptors according to sequences in the NCBI Genbank XM_015303261.1, XM_006352114.2 and XM_006354988.2. Primer sequences are shown in Supplementary Table S2. PCR products were gel purified and cloned using the PCR Cloning Kit (Thermo Scientific) into the plasmid pJET1.2/blunt according to the manufacturer’s instructions followed by transformation of *E. coli* strain DH10B (Invitrogen). *StHK4* was amplified using StHK4_truncated primers. The product was inserted into the construction of pB7FWG2-AHK3 instead of *AHK3*. The latter was removed at the BcuI and EcoRI restriction sites (Lomin *et al*., 2015). The nucleotide sequences of the cloned genes were confirmed by DNA sequencing.

*StHK2 and StHK3* sequences were subcloned into the plasmid pDONR™ 221 in BP reaction with Gateway® BP Clonase® II Enzyme mix (Thermo Scientific). Then, using the LR reaction with the LR Clonase® II Plus enzyme (Thermo Scientific), the cloned sequence was transferred into the expression vector pB7FWG2 (Karimi *et al*., 2007) where it was fused at the 3’-terminus to the *eGFP* gene. For expression in *E. coli*, *StHK2* and *StHK4* were amplified using primers StHK2_COLD and StHK4_COLD, respectively (Supplementary Table S2). The product was then inserted into the plasmid pCOLD IV (Takara BioInc.) at the XhoI and XbaI restriction sites for *StHK2* and SacI and EcoRI restriction sites for *StHK4*, followed by transformation of the *E. coli* DH10B strain.

### Transient expression of receptor genes in tobacco plants

The transient transformation of tobacco (*Nicotiana benthamiana* Domin) leaves was accomplished according to Sparkles *et al*. (2006). Eight week-old tobacco plants were infiltrated with a mixture of *Agrobacterium tumefaciens* carrying CK receptor genes fused to GFP and the *A. tumefaciens* helper strain p19 (Voinnet *et al*., 2003), and the expression of receptor genes was checked after 5–6 days on fluorescence microscope Axio Imager Z2 (Carl Zeiss Microscopy GmbH) before leaves were proceeded further for microsome isolation.

### Plant membrane isolation

All manipulations were done at 4 °C. Tobacco leaves 6 days after infiltration were homogenized in buffer (3 ml per 1 g of fresh weight) containing 100 mM Tris-HCl (pH 8.0), 2 mM Na_2_-EDTA, 50 mM KCl, 1 mM DTT and 1 mM PMSF. The homogenate was filtrated through Miracloth (Calbiochem, San Diego, USA), and the filtrate was centrifuged for 5 min at 5000 *g*. Then supernatant was centrifuged for 40 min at 15000 *g*. The microsome pellet was resuspended in 50 mM KCl, 10% glycerol and then microsome suspension was stored at −70 °C.

### Hormone binding assays

Ligand binding studies were performed in PBS as described previously (Romanov *et al*., 2005; Lomin *et al*., 2015). Studies of pH influence on hormone binding were performed in 50 mM MES-KOH (pH 5–7) or Tris-HCl (pH 7–9) buffers with 50 mM KCl. *K*_d_ for [^3^H]*t*Z binding to different receptors were determined in saturation assays followed by data analysis in Scatchard plots.

### Assessment of receptor functionality

Plasmids pCOLD IV with *StHK* coding sequences were transferred for the expression into *E. coli* strain KMI001 (Suzuki *et al*., 2001). In this strain, HK receptor→YojN→RcsB→*cps::lacZ* pathway can be activated by external CKs (Takeda *et al*., 2001). The activation of the signaling pathway was monitored by measuring β-galactosidase activity of *E. coli* cells. Cultivation of clones on Petri dishes containing 40 mM glucose, 40 μg ml^−1^ X-gal, 100 μM IPTG, 50 μg ml^−1^ ampicillin at 15 °C was performed for 4 days. The individual clones were then streaked onto new Petri dishes containing 40 mM glucose, 40 μg ml^−1^ X-gal, 100 μM IPTG, 50 μg ml^−1^ ampicillin ± *trans*-zeatin at a concentration of 0.5 μM. The clones were grown for 3 days at 15 °C. Expression of the *cps::lacZ* construct was evaluated by blue staining of bacterial clones.

### Gene expression analysis

Potato (*Solanum tuberosum* cv. Désirée) plants were cultivated under standard *in vitro* conditions at a long (16 h) day for 5-6 weeks on liquid MS medium containing 1.5% or 5% sucrose. For hormone treatment, the medium was replaced with the same one supplemented with *N*^6^-benzyladenine (BA, 1 μM). Tubes were inverted several times to assure uniform plant wetting and then incubated for 1 h under standard conditions. Finally plant organs (leaves, stems, roots, tubers) were isolated and immediately frozen in liquid nitrogen. Control plants were treated in a same way only without hormone. Total RNA was isolated by Trizol method (Brenner *et al*., 2005), this RNA served template for cDNA synthesis by reverse transcription (Invitrogen). All RNA samples were treated with RNase-free DNase I. The resulting cDNA was checked for the genomic DNA contamination by PCR with primers differentiating cDNA and genomic DNA. The band derived from genomic DNA was absent in the separating gel. Expression of genes encoding predicted proteins of CK signaling system was determined by qRT PCR. Potato housekeeping genes *StEF1*α (elongation factor 1-α, AB061263) and *StCYC* (cyclophilin, AF126551) were used as reference genes (Nicot *et al*., 2005). Sequences of primers for qRT PCR are shown in Supplementary Table S2.

### Statistical analysis

Statistical analysis was accomplished using the Student’s t-test. *P*-value <0.05 was considered as statistically significant. In tables and graphics, mean values with standard errors are presented.

## Results

### Monoploid Phureja genome analysis

#### Potato has everything necessary for CK signaling via the MSP pathway

The search for protein sequences and encoding genes involved in CK signaling was performed on the basis of the duplicated potato monoploid Phureja genome (Potato Genome Sequencing Consortium, 2011). In general, all potential components of the canonical CK signaling system described in Arabidopsis and other plant species with a sequenced genome (Kieber and Schaller, 2014; 2018) were identified in potato too. Potential CK-related genes found in potato encode homologs of CHASE domain-containing histidine kinases (CHK), phosphotransmitters (HPt), and response regulators of A (RR-A) and B (RR-B) types (Table 1). This indicates the MSP functioning in potato cells for CK signal transduction, involving proteins of a two-component system. In the potato monoploid proteome three predicted protein-coding sequences XP_015158747.1, XP_006352176.1 and XP_006355050.1, orthologous to Arabidopsis receptors AHK2, AHK3 and CRE1/AHK4, respectively, were detected. By analogy with the Arabidopsis orthologs, these proteins were annotated in NCBI as StHK2, StHK3, and StHK4. They correspond to mRNA sequences XM_015303261.1, XM_006352114.2 and XM_006354988.2. Deduced proteins StHK2, StHK3, and StHK4 share 59.35%, 67.75%, and 67.52% sequence similarity with the Arabidopsis orthologs. The lengths of *StHK2*, *StHK3* and *StHK4* genes are 5345, 4216, and 3810 bp, respectively, and predicted proteins are 1263, 1032, and 992 aa long (Table 1).

#### Phylogenetic analysis classified StHKs into three clades

The phylogenetic analysis was performed to compare the conserved and unique features of predicted potato CK receptors with the features of Arabidopsis, rice, tomato, and other species receptors (Fig. 1). CK receptors of flowering plants can be grouped into three main clades, corresponding to the Arabidopsis AHK2, AHK3, and CRE1/AHK4 receptors (Pils and Heyl, 2009; Lomin *et al*., 2012; Steklov *et al*., 2013). Predicted potato and tomato receptors are unequivocally distributed among these three clades. Evolutionally, they are closer to Arabidopsis than to rice receptors, what was expected since potato, tomato and Arabidopsis are dicots whereas rice is a monocot.

**Fig. 1.**
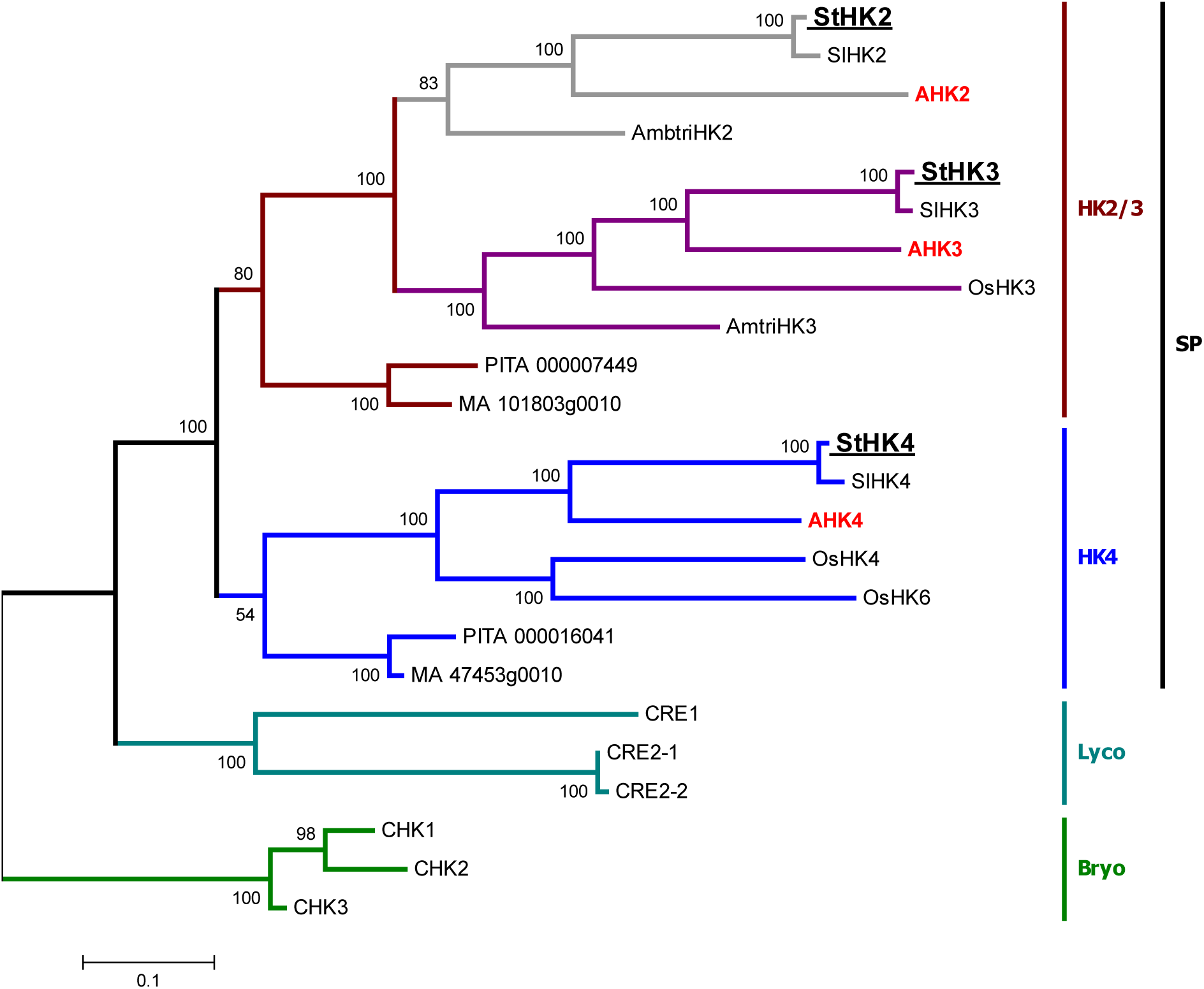
Phylogenetic tree of CK receptors. Species are: StHK2-4, *Solanum tuberosum*; SlHK2-4, *Solanum lycopersicum*; AHK2-4, *Arabidopsis thaliana*; OsHK3,4,6, *Oryza sativa*; AmbtriHK2,3, *Amborella trichopoda*; PITA 000007449 and PITA 000016046, *Pinus taeda*; MA 101803g0010 and MA 47453g0010, *Picea abies*; CRE1,2-1,2-2, *Selaginella moellendorffii*; CHK1-3, *Physcomitrella patens*. SP – seed plants, Lyco – Lycophyta, Bryo – Bryophyta. Parameters of ClustalW algorithm were: phylogeny test – bootstrap method, no. of bootstrap replications – 100, substitutions type – amino acid, model – equal input model, rates among sites – gamma distributed, no of discrete gamma categories – 3, gaps/missing data treatment – complete deletion, ML heuristic method – subtree-pruning – regrafting.

#### Multiple alignments revealed common and unique features of StHKs

We investigated the modular architecture of predicted potato CK receptors. The exon-intron structure of the cognate genes as well as occurrence and position of functional domains in the receptor proteins were analyzed. Known CK receptors share a common organization, including (from N to C termini) sensory module with CHASE domain, catalytic module with HisKA and ATPase domains, and receiver module with pseudoreceiver and receiver domains (Kakimoto, 2003; Steklov *et al*., 2013). The sensory module is flanked by predicted transmembrane (TM) α-helices. There is always a single TM-helix C-terminal (downstream) of module while the number of TM-helices N-terminal (upstream) of module is variable. Number of upstream TM-helices is usually highest (up to 3-4) in AHK2 clade members, lowest (1) in the AHK4 clade and intermediate in the AHK3 clade (Steklov *et al*., 2013). The domain structure of putative potato receptors fully corresponds to the canonical one (Fig. 2).

**Fig. 2.**
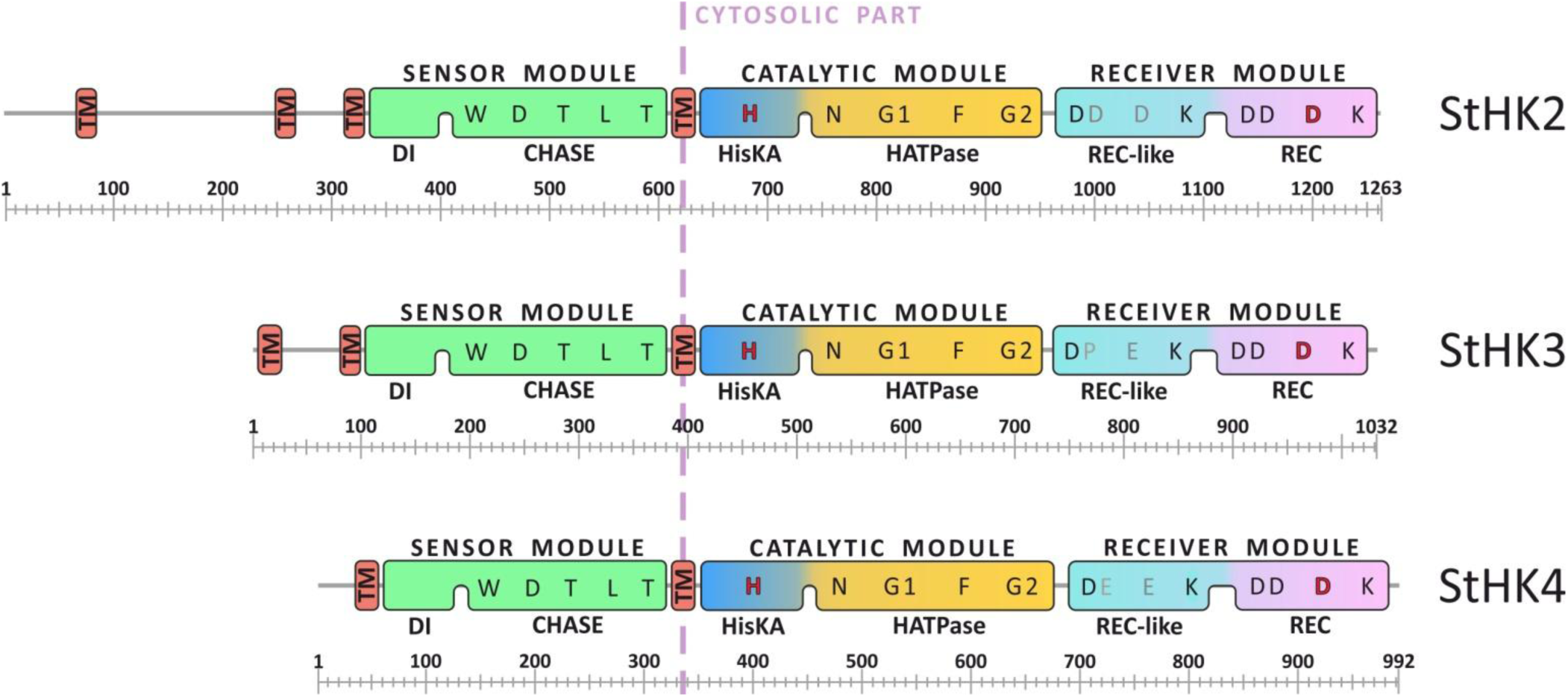
Module/domain structures of the predicted potato CK receptors. Protein domains: TM, transmembrane segment; DI, dimerization interface; CHASE, Cyclase/Histidine kinases Associated SEnsory domain (Steklov *et al*., 2013); HisKA, histidine kinase A domain; HATPase, adenosine triphosphatase domain; Rec-like, receiver-like domain; Rec, receiver domain. Conserved amino acids and consensus motifs (N, G1, F, G2) are indicated. According to conventional terminology, the catalytic module consists of dimerization and histidine phosphotransfer domain (DHpD), and catalytic and ATP-binding domain (CAD) (Mayerhofer *et al*., 2015; Pekárová *et al*., 2016). Scales at the bottom of the structures indicate the length in aa number.

At the N-termini of potato CK receptors, the number of upstream TM helices is 3, 2, and 1 in StHK2, StHK3, and StHK4, respectively. CK receptor genes share similar exon-intron organization. The exon boundaries in the receptor genes of different species coincide in most cases. A multiple alignment of receptor sequences from potato, rice and Arabidopsis was carried out (Fig. 3). All canonical motifs present in known CK receptors were also found in the potato orthologs. <u>H</u>, N, G1, F, and G2 motifs were identified in the catalytic module, and DD-<u>D</u>-K motifs – in the receiver domainof putative potato receptors. Conserved sequences contain phosphorylatable histidine (<u>H</u>) and aspartate (<u>D</u>) residues. StHK2 has a conserved aspartate in its receiver-like domain (Rec-like), similarly to orthologs from Arabidopsis (AHK2), tomato (SlHK2) and rice (OsHK3 and OsHK5). However, the overall DD-<u>D</u>-K-like motifs in Rec-like domains have little in common with the respective sequences in Rec domains (Fig. 3C).

**Fig. 3.**
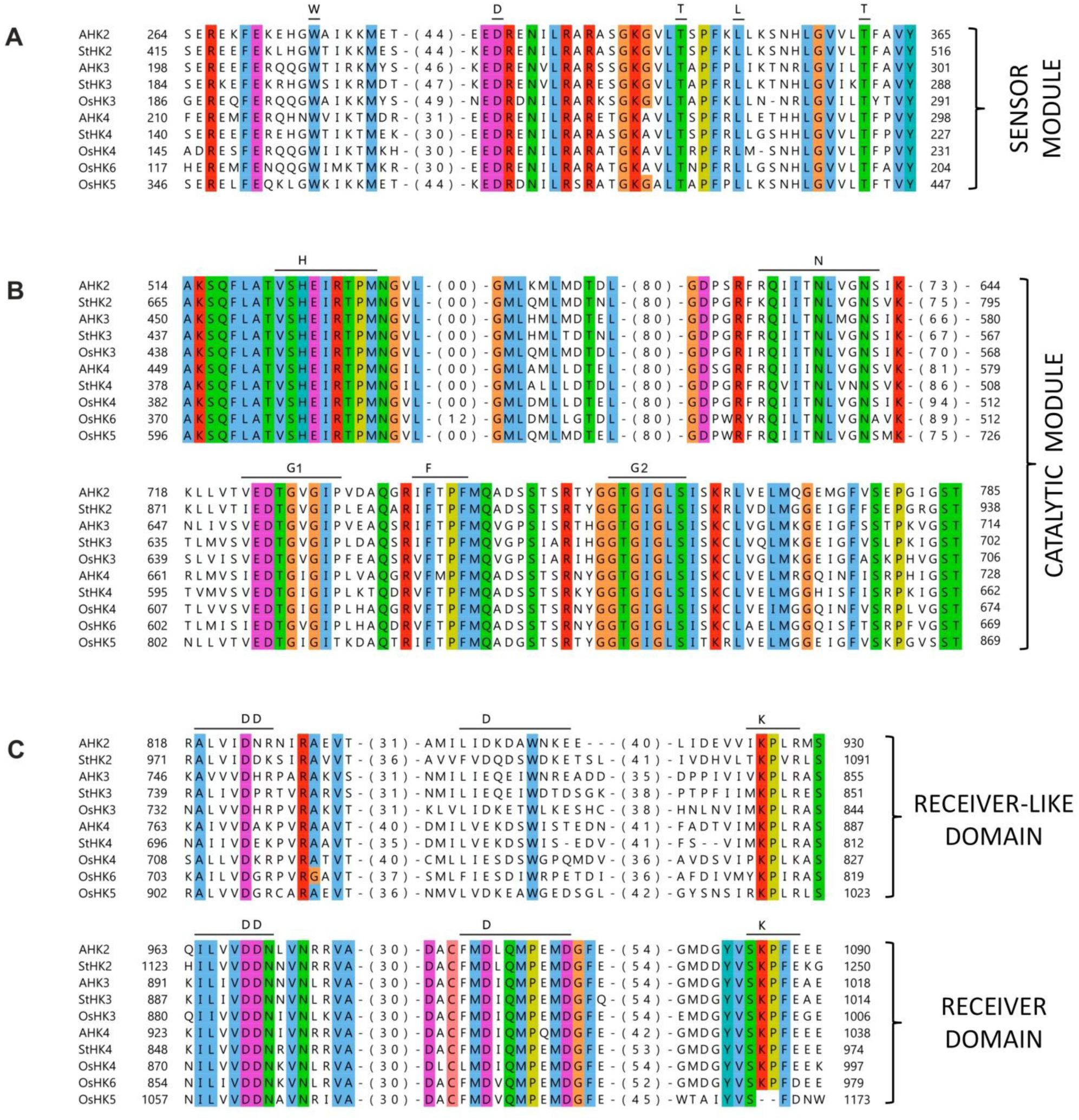
CK receptor sequence alignment. Consensus motifs and conserved aa are marked. AHK and OsHK correspond to Arabidopsis and rice proteins, respectively. Numbers of not shown aa are indicated in brackets.

Highly conserved motifs were earlier found in sensory modules and adjacent downstream TM-segments of CK receptors (Steklov *et al*., 2013). These motifs are obviously important for ligand binding and transmembrane signal transfer. In putative potato receptors, these motifs are also present, although with some peculiarities. In particular, StHK2 has a deviation from the canonic motif in CHASE domain, where either Glu or Asp is located at position 90, while StHK2 has Gln at this position. StHK3 has a deviation at the position 177, strongly conserved in the HK3 clade. This position is occupied by Phe in the canonic motif, while in StHK3 by Leu. In the general HK motif, either Phe or Tyr is located at position 177. StHK4 is distinguished by positions 83 (Ala→Ser) and 172 (Tyr→Phe) in conserved motifs. Note that counterparts of Gln90, Leu177 and Ser83 are present also in tomato genome, so these substitutions may be characteristic of Solanaceae family. Phe172 seems to be unique for potato.

#### StHK functional domains adopt canonical 3D structures

We have built homology models of all StHK domains (Fig. 4). High structural similarity of predicted potato receptors with their Arabidopsis orthologs was observed as expected. Key functional regions, such as ligand-binding sites, phosphorylation sites, ATP-binding sites and dimerization interfaces, are particularly conserved. Sensory modules consisting of dimerization, PAS and pseudo-PAS domains (the latter two comprise the CHASE domain) are very similar in Arabidopsis and potato. StHK2 and StHK3 differ from StHK4 by an insertion of 14 and 17 residues, respectively, in the region adjacent to the C-terminus of α3-helix (the first α-helix of the PAS domain). This insertion apparently does not participate in the hormone recognition site and is unlikely to directly affect the ligand-binding properties of the protein. Similar insertions are also present in AHK2 and AHK3 receptors from Arabidopsis.

**Fig. 4.**
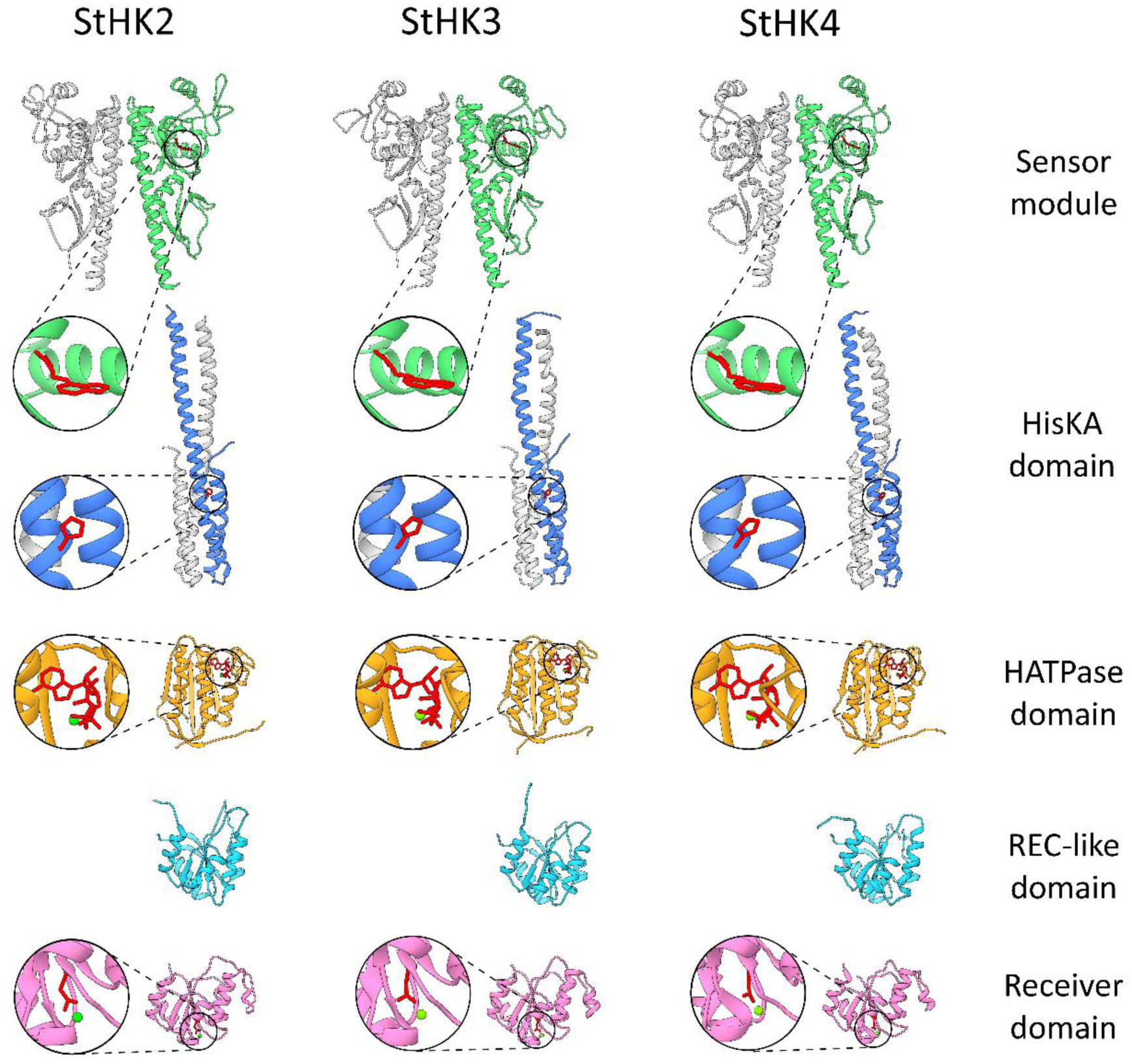
Homology models for predicted potato CK receptor domains. Sensor modules and HisKA domain are presented as dimers where one of subunits is colored grey. Positions of hormone, ATP and phosphoaccepting His/Asp residues are highlighted (red). Green spheres represent Mg^2+^ ions.

The catalytic modules include HisKA domains and H-ATPase domains. HisKA domains are formed by two α-helices and contain dimerization interface and phosphorylation site (conserved histidine). H-ATPase domains including ATP-binding sites have a sophisticated structure based on parallel/antiparallel β-strands and α-helices. A large insert at the β2-β3 linker (more than 50 residues long) differs CK receptors from bacterial histidine kinases and H-ATPase domain of the ethylene receptor. This insert is located, however, on the opposite side from the ATP binding site. This structural feature distinguishes not only potato receptors but also CK receptors of other species.

The CKI1 histidine kinase receiver domain (RD), used as the template for CK receptor RD, adopts a fold typical for the REC (or CheY-like) superfamily proteins. It is formed by five α-helices and β-sheet composed of five parallel β-strands. Two α-helices are located on one side of the β-sheet, and remaining three on the other side. The same fold is characteristic for the model of the Arabidopsis CRE1/AHK4 receptor RD. As distinct from this, an additional small helix is present in the region between α3 helix and β4 strand in the models of potato and other Arabidopsis receptors AHK2 and AHK3 RDs. Conserved aspartate residue, serving as a phosphate acceptor in RD, is located at the N terminus of the β3 sheet (Fig. 4).

Deviations from canonic CHASE motifs in sensory modules of putative potato CK receptors do not seem to alter 3D structures of the modules. Unusual Gln90 resides far from the ligand-binding pocket of StHK2, with sidechain directed to the dimerizing interface. Although the unusual Leu177 of StHK3 is localized in the ligand-binding site, its sidechain is oriented to the opposite direction. The substitutions in StHK4 seem to be more functional than in other predicted potato receptors. Ser73 and Phe172 are localized in the ligand-binding pocket periphery and their sidechains are oriented inwards. Hence, these latter substitutions might somehow influence the ligand specificity of the receptor.

### Experimental studies on autotetraploid potato cv. Désirée

#### Potato cv. Désirée possesses multiple alleles of StHK genes

A homozygous doubled monoploid Phureja (DM1-3 516 R44) is an artificial form of potato phenotypically differing from commonly known diploid/tetraploid potato varieties (Potato Genome Sequencing Consortium, 2011). Such differences in phenotype are underlain by considerable sequence and structural genome variations between potato haplotypes. Therefore, the results of genome study of monoploid Phureja do not mirror exactly more complex genomes of common potato cultivars.

Our experimental study of CK receptors was performed using the autotetraploid potato cv. Désirée, widely used for commercial and scientific purposes (Aksenova *et al*., 2000; Kolachevskaya *et al*., 2015). We cloned the putative receptor genes using primers designed according to Phureja gene sequence data. Distinct from Phureja genome, at least six genes of putative CK receptors were cloned from cDNA of Désirée plants. All these genes share a typical module/domain structure characteristic of hybrid sensor histidine kinases (Figs. 2–4). According to their sequence, encoded proteins fall pairwise into three known clades of CK receptors (Table 2, Fig. 1). Thus, each form of CK receptors from potato cv. Désirée consists of at least two close isoforms encoded by natural receptor alleles. Sequencing of cloned genes revealed traits of both similarity and divergence between Phureja and Désirée plants. The nucleotide sequences of HK2-clade members *StHK2a* and *StHK2b* differ from the orthologous Phureja sequence by five and four nucleotides (5 and 4 SNPs), respectively. At the protein level, StHK2a and StHK2b have three and two aa substitutions, respectively, relative to Phureja receptor (Table 2).

**Table 2.** Putative CK receptor genes in potato genomes and encoded proteins

| Receptor clade | Putative CK receptors of potato plants: |  | SNP number in putative CK receptor genes/proteins of cv. Désirée vs var. Phureja: |  | Number of Désirée cDNA clones |
| --- | --- | --- | --- | --- | --- |
|  | Phureja*<br>(length, aa) | Désirée **<br>(length, aa) | DNA bases | Amino acids |  |
| HK2 orthologs | StHK2<br>(1263 aa) | StHK2a (1263 aa) | 5 SNPs | 3 SNPs | 17 |
|  |  | StHK2b (1263 aa) | 4 SNPs | 2 SNPs | 17 |
| HK3 orthologs | StHK3<br>(1032 aa) | StHK3a (1032 aa) | No SNP | No SNP | 6 |
|  |  | StHK3b (1031 aa) | 20 SNPs, 3 del. | 9 SNPs, 1 del. | 3 |
| CRE1/AHK4 orthologs | StHK4<br>(992 aa) | StHK4a (992 aa) | No SNP | No SNP | 1 |
|  |  | StHK4b (991 aa) | 28 SNPs, 3 del. | 13 SNPs, 1 del. | 9 |

\*Doubled monoploid, method: total genome sequencing.
\*\* Autotetraploid, method: PCR with cDNA as a template.

Of two cloned genes of HK3-clade, *StHK3a* is identical to its counterpart of Phureja, whereas *StHK3b* differs by 20 SNPs together with a 3-nucleotide deletion. These differences result in the absence of one aa and nine aa substitutions in StHK3b compared to its Phureja ortholog. Similar data were obtained for HK4-clade: *StHK4a* was fully identical to that of Phureja whereas *StHK4b* differs by 28 SNPs and a 3-nucleotide deletion. Correspondingly, StHK4b differs from its Phureja ortholog, as well as from StHK4a, by deletion of one aa and substitution of 13 ones (Table 2). Analysis of aa sequences of the proteins showed that all putative histidine kinases of Désirée potato retain the domains and consensus sequences typical for CK receptors, despite aa substitutions (Fig. 2). This indicates that all proteins encoded by the cloned *StHK* genes of tetraploid potato plants can successfully function as CK receptors.

#### StHKs have typical CK-binding properties except StHK3 with distinct ligand specificity

To analyze ligand-binding properties of the receptors, a recently developed plant membrane assay system (Lomin *et al*., 2015) was used. Predicted potato CK receptor genes were cloned into pB7FWG2 vectors for transient expression in tobacco leaves. In the case of *StHK2* and *StHK4* genes, the full-length cDNA sequences were expressed, but in the case of *StHK3*, expression of the full-length receptor failed for unknown reasons. Instead of full-length receptor, we cloned a genomic sequence of the StHK3a sensory module flanked with transmembrane domains. From the transiently transformed tobacco leaves, a microsomal fraction enriched with individual potato receptors was obtained. The binding assays were conducted using this fraction and tritium-labeled CK. In aggregate, we tested four putative receptors belonging to all three clades: StHK2a, StHK3a (sensory module, further designated as StHK3a_SM_), StHK4a, and StHK4b.

First, we determined the pH-dependence of hormone binding to these receptors within the pH range of 5–9 (Fig. 5). All StHKs exhibited maximal *trans*-zeatin binding at the neutral-mildly basic pH: StHK2a at pH 7.5, StHK3a_SM_ at pH 7, StHK4a at pH 7.5–8, and StHK4b at pH 8–9. All StHKs showed a decrease in ligand binding at acid pH: StHK2a and StHK3a_SM_ reduced their binding at pH 5 compared to pH 7 by a factor of 2 and 5, respectively. Ligand binding by StHK4a and StHK4b decreased at pH 5 about three times compared to maximal values. Although the StHK3a was represented in this study only by its sensory module, a control experiment with the full-length StHK2a and its sensory module showed a similar pH-dependence of hormone binding (data not shown). This means that an isolated sensory module is sufficient to determine the pH-dependence of hormone binding by the receptor.

**Fig. 5.**
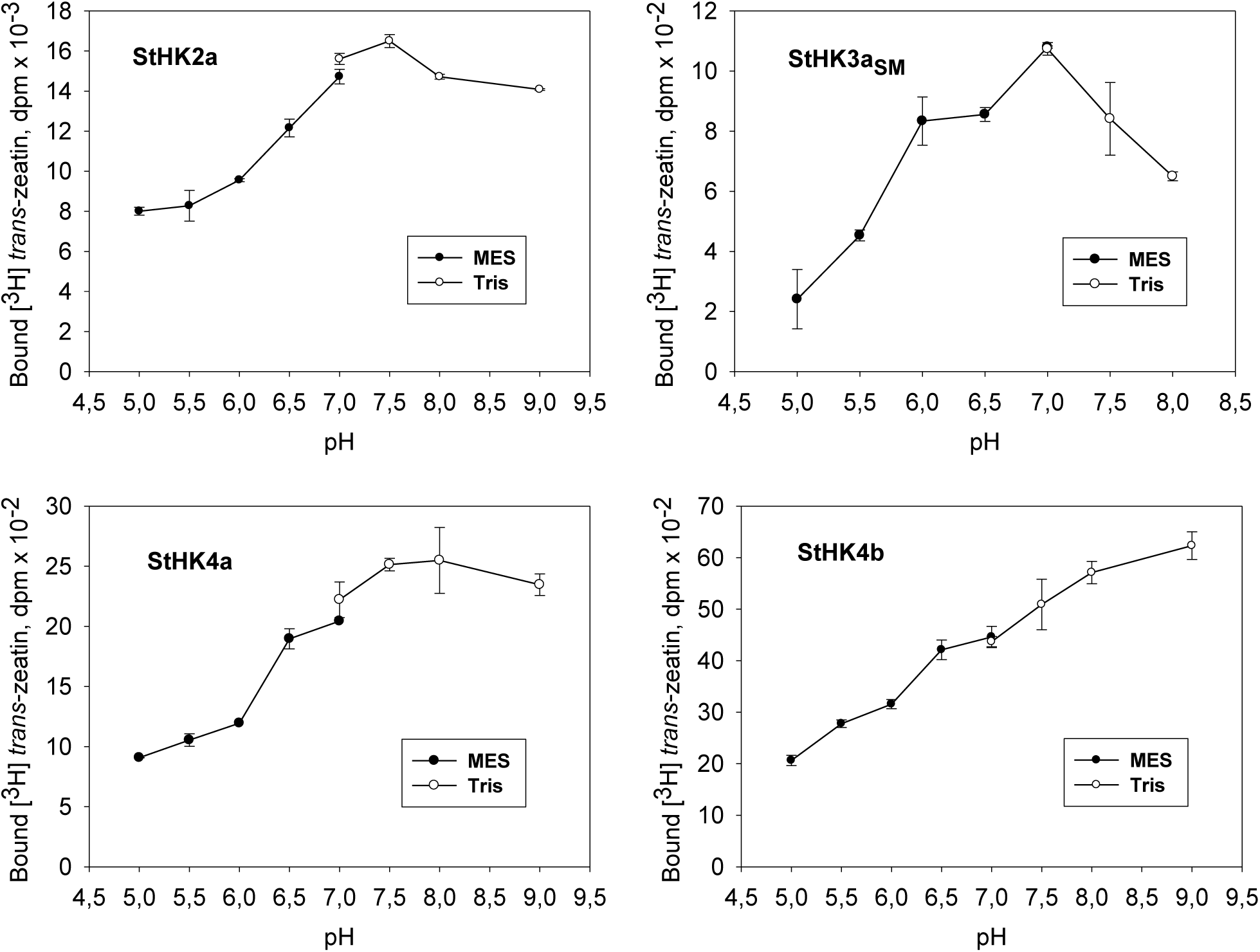
pH dependencies of *trans*-zeatin binding to putative potato CK receptors.

The interaction of a hormone with a receptor is characterized by the equilibrium dissociation constant (*K*_d_) of the ligand-receptor complex. *K*_d_ values were determined by the dose-dependent binding of labeled *trans*-zeatin to StHKs, the results were processed by the Scatchard method (Supplementary Fig. S7) (Lomin and Romanov, 2008). All StHKs demonstrated high affinity for *trans*-zeatin, with similar *K*_d_ at the nanomolar level (Table 3). The determined *K*_d_ values were close to the values of analogous constants for CK receptors of other species (Lomin *et al*., 2012, 2015, Kuderová *et al*., 2015) and were well correlated with concentrations of active CKs *in planta* (Hirose *et al*., 2008) including potato (Kolachevskaya *et al*., 2017, 2018).

**Table 3.**
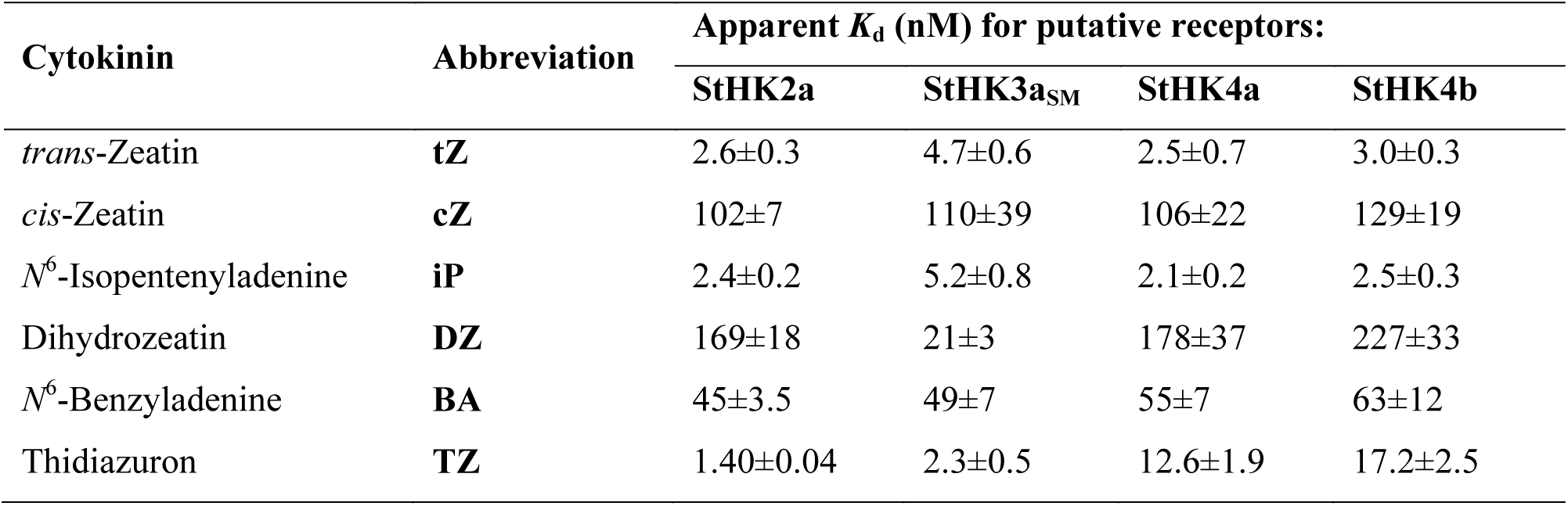
The affinity (K_d_) of various CKs for putative potato receptors

Different CKs are usually present in the plant: *trans*- and *cis-*zeatins, isopentenyladenine, and dihydrozeatin. In addition to natural CKs, there are many synthetic ones. Receptors exhibit different affinities for these compounds (Lomin *et al*., 2015; Savelieva *et al*., 2018). We studied the ligand specificity of putative receptors in competitive experiments where binding of labeled CK was carried out in the presence of various concentrations of certain unlabeled ligands. Based on the obtained competition curves, the apparent *K*_d_ values were determined for each ligand as described (Lomin and Romanov, 2008). We analyzed the interaction of StHKs with six CKs, including five natural ones as well as synthetic urea-type CK thidiazuron (Table 3). The ligand specificity of StHKs showed much in common. All analyzed proteins had a high and nearly equal affinity for *trans*-zeatin and isopentenyladenine, apparent *K*_d_ ranging from 2.1 to 5.2 nM. All StHKs bound *cis*-zeatin significantly weaker, with *K*_d_ over 100 nM. *N*^6^-Benzyladenine exhibited an intermediate affinity with *K*_d_ ranging from 40 to 60 nM. Regarding the two remaining CKs, StHK proteins showed significant differences. StHK3a_SM_ bound dihydrozeatin with *K*_d_ ~21 nM, much stronger than other putative potato receptors (*K*_d_~170-230 nM). StHK2 and StHK3a_SM_ showed a high affinity for thidiazuron (*K*_d_=1.4 nM and 2.3 nM, respectively), whereas its affinity for StHK4a and StHK4b was much lower (*K*_d_=12.6 nM and 17.2 nM, respectively). The CK affinity ranking for StHKs was as follows: StHK2, TD>iP=tZ>BA>cZ>DZ; StHK3, TD>iP=tZ>DZ>BA>cZ; StHK4, iP=tZ>TD>BA>cZ>DZ. The preference profiles of StHK2 and StHK3a_SM_ differ by DZ position, and from (almost identical) StHK4 isoforms by TD position. The greatest differences (in TD and DZ positions) were revealed between StHK3 and StHK4. Although StHK3a was represented in this study only by its sensory module, previous data showed that sensory module is sufficient to characterize the ligand preference of the full-length receptor (Stolz *et al*., 2011; Lomin *et al*., 2015).

#### StHKs are able to trigger signaling via MSP

The ability of the putative potato receptors to trigger CK signaling was tested on *E. coli* ΔrcsC mutant devoid of its own RcsC hybrid histidine kinase and equipped with the *cps:LacZ* construct with the *LacZ* reporter gene driven by *cps* promoter (Suzuki *et al*., 2001; Takeda *et al*., 2001). This design allows assessment of the ability of hybrid histidine kinases to initiate signaling over the MSP pathway. Activation of MSP signaling in the bacteria leads to the expression of the reporter galactosidase (LacZ), whose activity is manifested by blueing of clones growing on X-Gal-supplemented medium. We expressed the cloned genes of the putative potato CK receptors in *E. coli* ΔrcsC. In the clones expressing the StHKs but not in the control clone, blue staining was observed (Fig. 6). The degree of blueing was greatly increased in the presence of CK. It confirms the ability of the cloned potato proteins to transmit the CK signal to the primary response genes via the canonic MSP pathway.

**Fig. 6.**
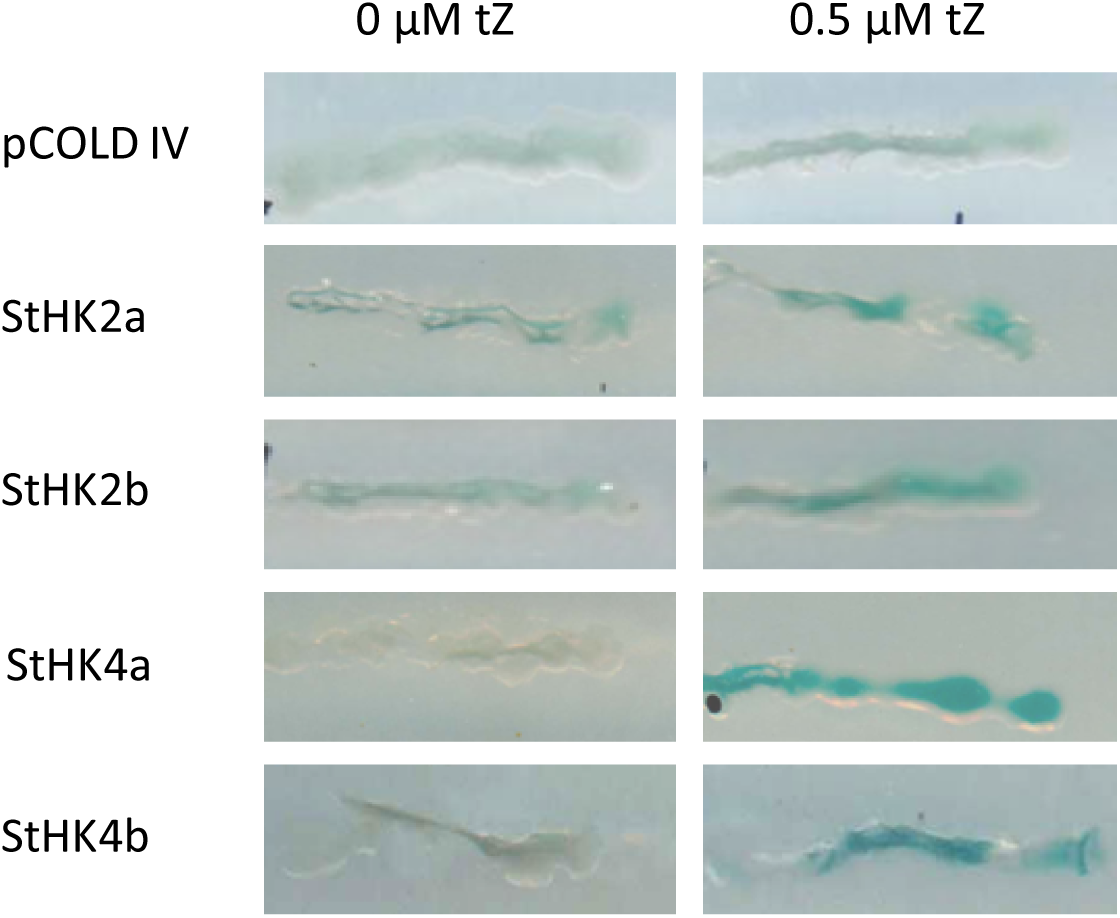
CK receptors of potato feed MSP signaling pathway in ΔRcsC *E. coli* cells.

#### StHKs exhibit *in planta* organ-specific expression pattern which has unique properties

To assess the functionality of a gene *in vivo*, it is important to know the level and pattern of its expression in the living organism. We studied the expression of putative CK receptor genes in organs of potato plants grown *in vitro* under conditions favorable for either vegetative growth (1.5% sucrose) or for tuber formation (5% sucrose). The mRNA contents of the *StHK2*, *StHK3* and *StHK4* genes was determined by the qRT PCR method. For the quantitative comparison of the expression profiles, intra-exon primers were selected for each tested gene (Supplementary Table S1). These primers were complementary to both alleles of the same clade owing to a great similarity of these gene sequences. The relative amounts of putative receptors of distinct clades in potato organs were judged by comparing the levels of transcripts of the cognate genes.

Expression levels differed significantly depending on *StHK* group, organ and growth conditions (Fig. 7). Expression patterns were different in plants grown on media with low (1.5%) or high (5%) sucrose content. In the case of 1.5% sucrose medium, the highest expression of *StHK3* genes was observed in roots, while in the case of 5% sucrose medium, the maximal expression of *StHK3* tended to occur in leaves. In the low sucrose grown plants, *StHK4* gene was much weaker expressed in leaves than in stems or roots, whereas at the higher sucrose content levels of *StHK4* expression in different organs were more equalized. In the *StHK2* group, noticeable organ-specific differences were detected when plants were grown on 5%, but not on 1.5% sucrose. The lowest expression level of all StHK groups was usually observed in tubers compared to other organs (Fig. 7A).

**Fig. 7.**
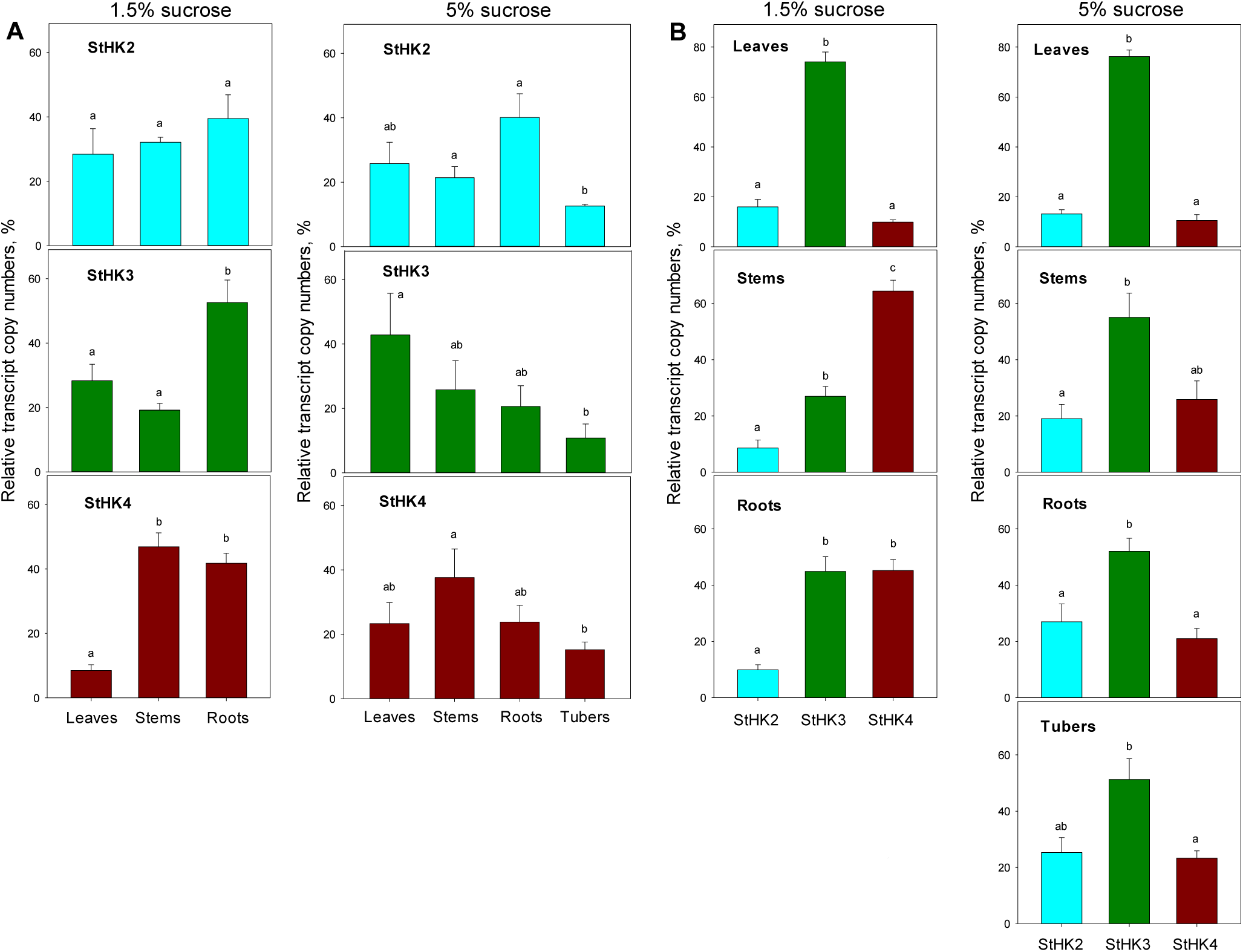
Organ-dependent (A) and clade-dependent (B) patterns of expression of CK receptors in potato plants cultivated on media with different % sucrose. Relative transcript copy number is given as % of the total transcript amount in each plot, taken as 100%. Different letters (a, b, c) indicate significant differences at *P* < 0.05.

Within each organ, expression of *StHK3* undoubtedly dominated in leaves, regardless of the sucrose content (Fig. 7B). Expression of *StHK2* and especially *StHK4* genes in leaves was much weaker. In stems grown on 1.5% sucrose, expression of *StHK4* prevailed, while the lowest expression was characteristic of *StHK2* genes. In the roots, expression of *StHK2* genes was relatively weak, whereas the genes of *StHK3* and *StHK4* clades were expressed actively and in almost equal proportions. A dissimilar pattern of expression was observed in plants grown on 5% sucrose. Here in addition to leaves, in all other organs tested (stems, roots, tubers) the expression of *StHK3* alleles prevailed too, though to a lesser extent than in leaves. Compared to the low-sucrose medium, 5% sucrose increased the relative expression of *StHK2* genes (in stems and roots), while decreasing the level of *StHK4* expression. Thus, unlike Arabidopsis, in potato plants there is evidently no dominance of StHK4 receptors in roots, on the contrary, StHK3 receptors seem to dominate there when cultivating plants on tuber-inductive 5% sucrose. A common feature of potato and Arabidopsis is a very low expression of HK4 orthologs in leaves.

Although the primers used for qRT PCR did not distinguish closely related isoforms of the CK receptor genes, it is still possible to approximately estimate the relative expression of these alleles. To achieve this goal, data on cDNA clone numbers can be used (Table 2). Within the same clade, relative quantity of cDNA clones harboring a distinct isoform should reflect the relative occurrence of cognate mRNAs. According to the last column of Table 2 corresponding to aerial part of potato seedlings, two mRNA isoforms of the HK2 clade were in the 1:1 ratio; among mRNA isoforms of HK3 clade, *StHK3a* was approx. two-fold more frequent than *StHK3b*; in the case of HK4 clade, *StHK4b* was expressed about one order of magnitude more intensively than *StHK4a*.

#### StHK promoter activity is hardly affected by CKs, in accordance with low *cis-*element content

Treatment of potato plants with *N*^6^-benzyladenine had a small effect on the expression of the CK receptor genes, and the hormonal impact, when occurred, was only local and not always reproduced. At 1.5% sucrose, the upregulation (on average, 2.5-fold) of *StHK4* expression was regularly recorded, but only in leaves (Fig. 8). It can be stated that the CK effect on the expression of potato receptor genes, if any, is mostly limited to *StHK4* and depends on both organ/tissue type and conditions of plant cultivation.

**Fig. 8.**
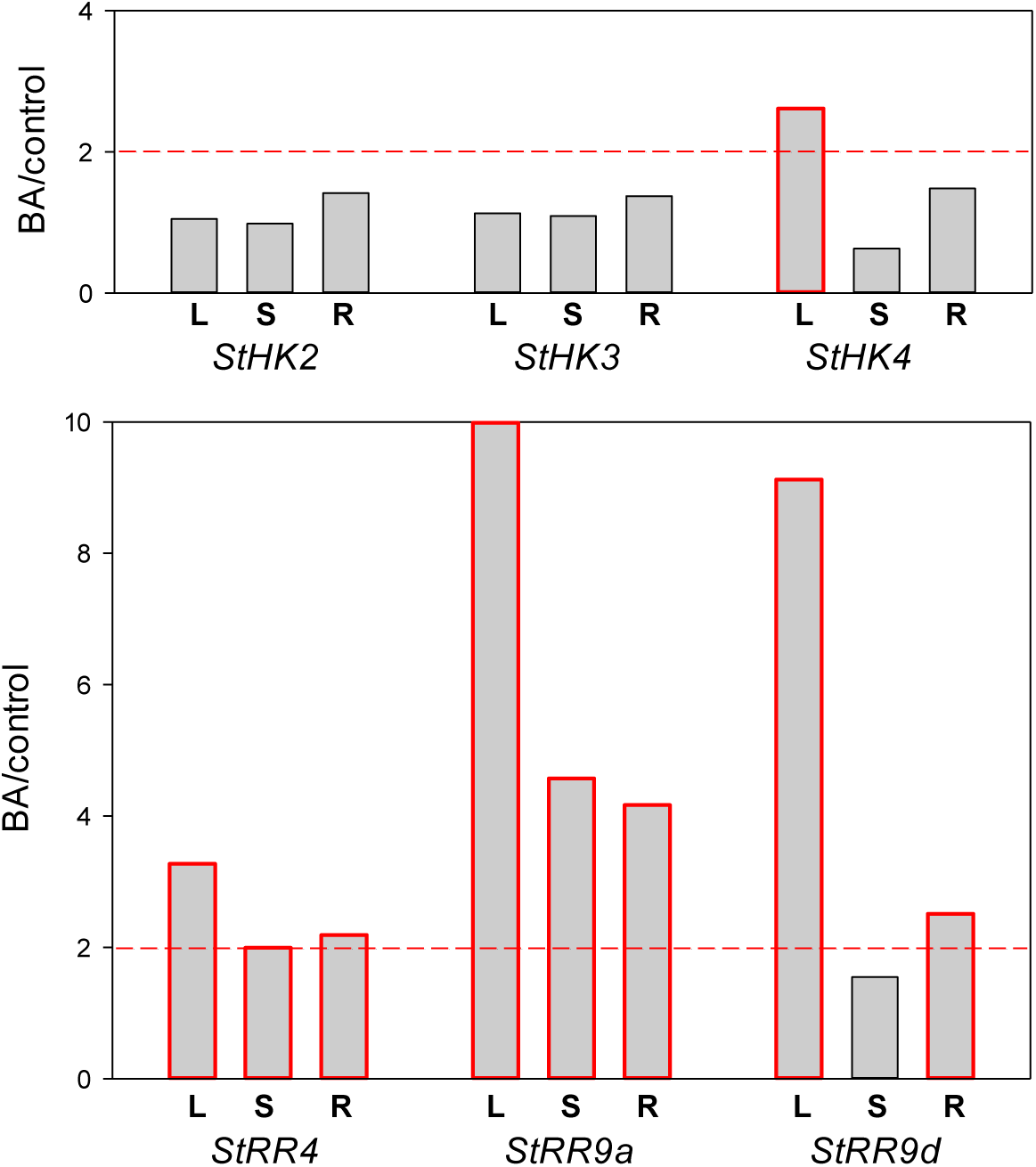
Degrees of transcription induction (BA/control) of CK receptor (top) and response regulator type A (bottom) genes after 1 h treatment of potato plants with 1 μM BA. Plants were grown on MS medium with 1.5% sucrose for 5-6 weeks under standard LD conditions. L, S, R signify leaves, stems and roots, respectively. More than two-fold prevalence of transcripts in BA-treated over control plants is considered as significant induction, bars corresponding to induced genes are outlined red.

To validate the results of CK treatment experiments, the effect of CK administration on the transcript level of the genes of type A response regulator (*RR*-A) genes was analyzed. These genes in other species (Arabidopsis, maize) represent genes of primary response to CK, so it might be expected that in potato too they would be responsive to CK. Indeed, our experiments showed a rapid and reliable increase in the expression of *StRR*-A genes, in contrast to the receptor genes, after plant treatment with BA (Fig. 8). These results prove the reliability of design and implementation of experiments and, on the other hand, corroborate the common mode of functioning of the CK signaling system in different plant species.

Analysis of promoter structures of the studied genes (Fig. 9) was mostly consistent with the gene expression data. Long CK-sensitive *cis*-regulatory elements or blocks of 4 or more short elements near the transcription start (~300 bp area) were found in promoters of almost all *StRR*-A, but not *StHK* genes. Among the receptor genes, only *StHK4* has a block of 3 short CK-sensitive *cis*-elements near the start of transcription. It is possible that this block determines the responsiveness of *StHK4* to CK under certain conditions, as shown in Fig. 8. Though this promoter analysis was accomplished using the genome sequence of var. Phureja, the promoter sequencing from Désirée plants showed an identity of the promoters from these two potato lines.

**Fig. 9.**
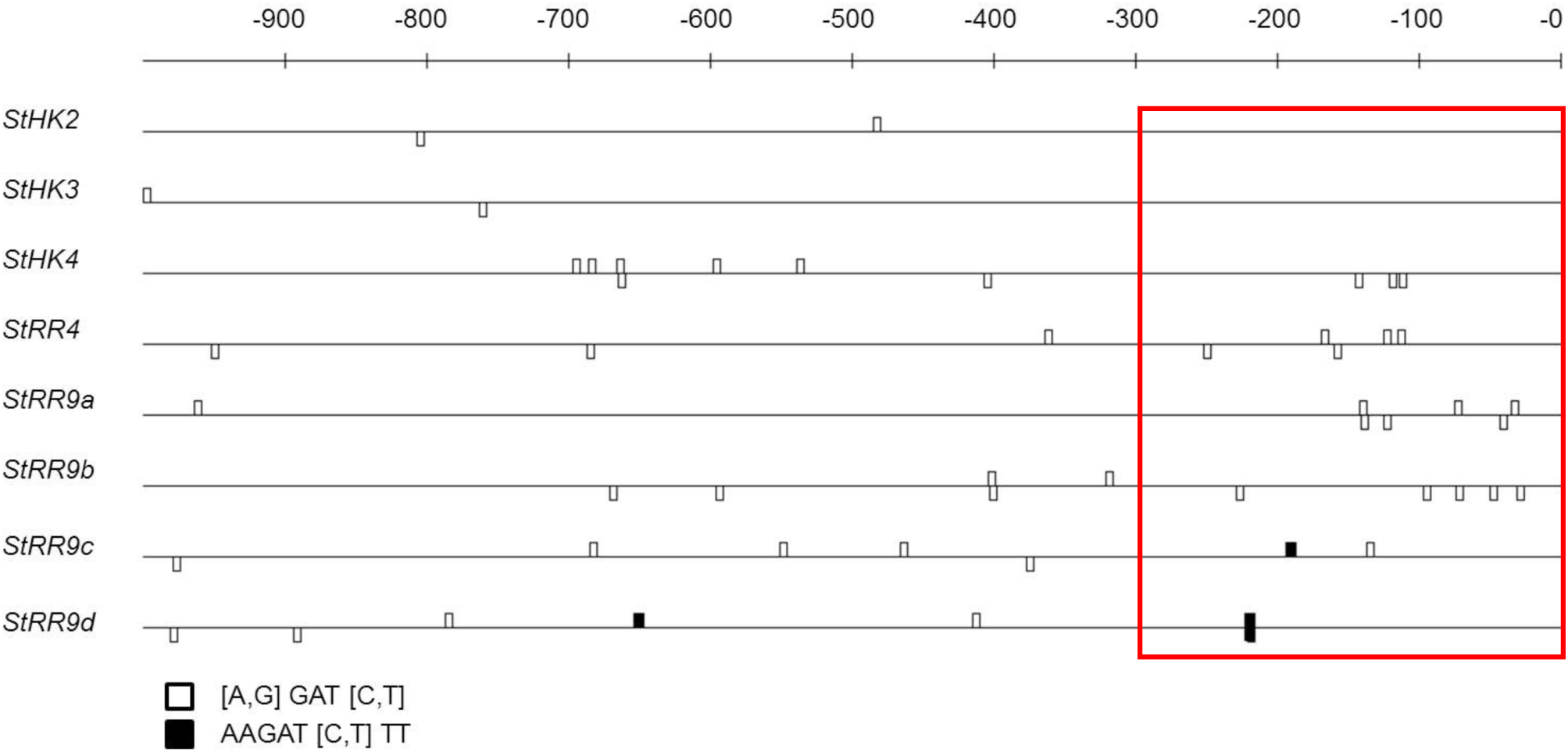
CK-responsive *cis*-regulatory elements in promoters of CK receptor genes (upper part) and response regulators type A genes (lower part) of potato. Elements are shown on both DNA strands. Promoter area proximal to transcription start is boxed.

## Discussion

Plant morphogenesis, in particular tuberization, is based on spatiotemporal cell proliferation and differentiation. The main biological effect of plant hormones CKs is the induction of cell divisions (Sakakibara, 2006; Romanov, 2009), therefore CKs are important participants of morphogenetic processes. Indeed, with regard to potato development, CKs were reported to accelerate and scale up tuber formation (Aksenova *et al*., 2000; Romanov *et al*., 2000). In non-potato plants, CKs alone were able to induce the emergence of tuber-like structures (Guivarc’h *et al*., 2002; Eviatar-Ribak *et al*., 2013; Frugier *et al*., 2008; Miri *et al*., 2016). Apart the impact on the formation of tubers, CKs are known to regulate overall plant architecture, biomass partitioning as well as resistance to biotic and abiotic stress-factors (Aksenova *et al*., 2000; Abelenda and Prat, 2013; Zwack and Rashotte, 2015; Brütting *et al*., 2017; Thu *et al*., 2017). All these point to the importance to investigate CK signaling system in plants, in particular in tuber crops like potato.

Herein, we present first results of detailed study of CK receptors from potato plants. Two different potato forms were examined: doubled monoploid Phureja and tetraploid potato of Désirée variety. Phureja plants possess, like Arabidopsis, three CK receptor orthologs. By contrast, in Désirée plants two allelic forms of each receptor type (StHK2a/b, StHK3a/b and StHK4a/b) have been found belonging to the three known phylogenetic clades. Our data indicated that this receptor abundance is characteristic of each individual Désirée plant. It is not excluded that the real number of receptor alleles in potato plant is somewhat higher. Within each group, receptor isomers differ by a few aa substitutions which do not affect most conserved motifs. However, some consensus motifs in sensory module (Steklov *et al*., 2013) are distinctive in receptors of potato. The reason for such peculiar properties is not yet clear. Molecular modeling was employed to build models of the structure for all main domains of potato CK receptors. In general, potato CK receptors share similar domain structure with crystallized hybrid histidine kinases from other species. Note that such a complete characterization of all main domains of CK receptors is presented for the first time.

The ligand-binding properties of individual potato receptors have been determined: affinity constants for active CKs, pH-dependence of ligand binding, ligand specificity. Two of the studied receptors (StHK3a and StHK4a) are identical in potato cv. Désirée and var. Phureja. All receptors have high affinity for tZ, significantly lower for BA, and relatively low for cZ. StHK3 differs from other potato receptors by relatively high affinity for DZ. The ligand specificity of StHK2 and StHK4 has much in common with that of Arabidopsis orthologs, whereas StHK3 binds iP and BA much strongert than AHK3, the affinity of StHK3 for iP and tZ is similar. Thus, the ligand-binding properties of StHK3 differ from those of orthologs in Arabidopsis, maize and oilseed rape. All receptors bind CK stronger in basic (pH 7–9) than acidic (pH 5–7) pH range. This evidences in favor of the intracellular functioning of potato CK receptors (Romanov *et al*., 2018). The functionality of cloned potato receptors was confirmed by testing their ability to transduce CK signal via MSP up to the target gene.

The predominant expression of the *StHK3* genes was revealed in leaves, as well as in other organs of plants grown on 5% sucrose, although the degree of dominance of StHK3 was less pronounced in stems, roots and tubers. When plants were grown on 1.5% sucrose, *StHK4* expression predominated in stems while in roots the expression levels of *StHK3* and *StHK4* were relatively high and nearly equal. In contrast to other species (Romanov, 2009; Lomin *et al*., 2012), no prevalent expression of HK4 orthologs in roots was found. Exogenous CK had little effect on the expression of CK receptors in potato plants except *StHK4* which can be rapidly upregulated in leaves. Analysis of promoter structures showed a correlation between the occurrence of *cis*-regulatory elements and the CK sensitivity of gene expression.

Thus, the totality of our results left no doubt that studied StHK proteins are genuine CK receptors in potato. The observed unique structural features refine and broaden our notion on the properties of CK receptors. The revealed peculiarities of CK perception apparatus in potato might be associated with the ability of this crop to produce tubers. It may be suggested that tuber initiation can be associated with the local/temporary increase in CK signaling in stolon tips. The obtained results create a solid basis for further in-depth study of the role of the CK signaling system in potato ontogenesis and provide new biotechnological tools to optimize hormonal regulation of tuber formation.

## Supplementary Data

**Table S1.** Sequence identity of modeled receptor domains and corresponding templates.

| Domain | Template |  | Receptor | Identity,<br>% | Reference |
| --- | --- | --- | --- | --- | --- |
|  | PDB ID | Protein |  |  |  |
| Sensory module | 3T4L_A | AHK4 | StHK2 | 64.75 | Hothorn *et al.*, 2011 |
|  |  |  | StHK3 | 65.00 |  |
|  |  |  | StHK4 | 79.62 |  |
| HisKA domain | 4MT8_A | ERS1 | StHK2 | 36.99 | Mayerhofer *et al.*, 2015 |
|  |  |  | StHK3 | 34.72 |  |
|  |  |  | StHK4 | 38.20 |  |
| ATPase domain | 4PL9_A | ETR1 | StHK2 | 34.94 | Mayerhofer *et al.*, 2015 |
|  |  |  | StHK3 | 32.74 |  |
|  |  |  | StHK4 | 31.55 |  |
|  | 5IDM_A | CckA | StHK2 | 19.88 | Dubey *et al.*, 2016 |
|  |  |  | StHK3 | 22.42 |  |
|  |  |  | StHK4 | 22.70 |  |
| Receiver domain | 3MMN_A | CKI1 | StHK2 | 48.21 | Pekárová *et al.*, 2011 |
|  |  |  | StHK3 | 43.64 |  |
|  |  |  | StHK4 | 48.78 |  |
|  | 4EUK_A | AHK5/CKI2 | StHK2 | 32.52 | Bauer *et al.*, 2013 |
|  |  |  | StHK3 | 31.88 |  |
|  |  |  | StHK4 | 32.37 |  |
| Receiver-like domain | 1DCF_A | ETR1 | StHK2 | 25.00 | Müller-Dieckmann *et al.*, 1999 |
|  |  |  | StHK3 | 15.45 |  |
|  |  |  | StHK4 | 21.77 |  |

**Table S2.**
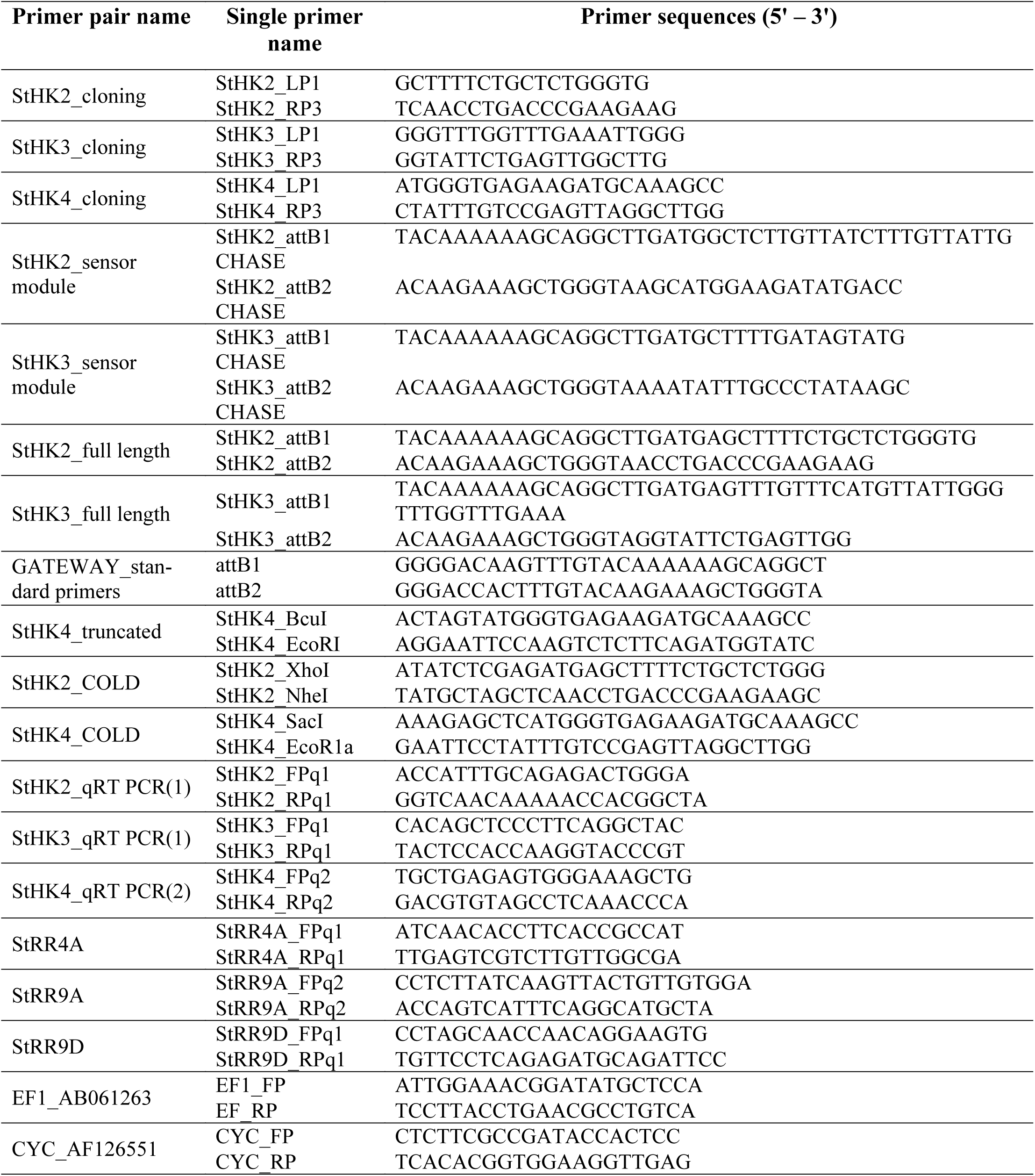
Primers used in this work.

## Acknowledgements

This work was supported by the Russian Science Foundation, grants no. 14-14-01095 (before 31.12 2016, bioinformatic and initial experimental data) and 17-74-20181 (in 2017, conclusive experimental results). We thank T. Schmülling for providing opportunity to perform some experiments in his laboratory.

**Abbreviations**
aa: amino acid
BA: 6-benzyladenine
CHASE: Cyclases/Histidine kinases Associated SEnsory
CHK: CHASE domain-containing histidine kinases
CK: cytokinin
CRF: cytokinin response factor
cZ: *cis*-zeatin
DI: dimerization interface
DZ: dihydrozeatin
GFP: green fluorescent protein
HK: histidine kinase
iP: isopentenyladenine
LacZ: galactosidase
LD: long day
MSP: multistep phosphorelay
RR: response regulator
SNP: single nucleotide polymorphism
TD: thidiazuron
TM: transmembrane
tZ: *trans*-zeatin.

